# Phylogeny, systematics and evolution of mimicry patterns in Neotropical limenitidine butterflies

**DOI:** 10.1101/2025.07.03.662938

**Authors:** V Erika Páez, Gael J. Kergoat, Nicolas Chazot, Mohamed Benmesbah, Adriana D. Briscoe, Susan D. Finkbeiner, André V.L. Freitas, Robert P. Guralnick, Ryan I. Hill, Marcus R. Kronforst, Luiza Moraes Magaldi, Sean P. Mullen, Ichiro Nakamura, Hannah L. Owens, Niklas Wahlberg, Maxwell Woodbury, Marianne Elias, Keith R. Willmott

## Abstract

The Neotropical butterfly genus *Adelpha* Hübner exhibits remarkable species diversity and striking convergence in wing colour patterns potentially explained by mimicry, making it an exceptional model for exploring trait evolution and its relationship with speciation. To date, unresolved phylogenetic relationships hinder a comprehensive understanding of the evolutionary biology of the genus. Using a novel multi-marker dataset combining one mitochondrial and 15 nuclear gene fragments, we generate the most comprehensive phylogeny of the genus *Adelpha* to revisit its systematics and investigate the evolution of mimicry colour patterns. Our data set encompasses 83 of the 87 known extant species and six *Limenitis* species that were recently excluded from *Adelpha* (134 of *c*. 160 subspecies in total), collectively displaying 14 distinct mimicry patterns. We provide conclusive evidence that corroborates previous work on the polyphyly of *Adelpha* as historically conceived, and describe the genus *Adelphina* Páez & Willmott **n. gen.** to stabilize the nomenclature, both genera representing Neotropical limenitidines. The comprehensive phylogeny provided in this study lays a solid foundation for future research into the processes driving diversification within these species interacting through mimicry. Ancestral character state reconstruction reveals gradual evolution of mimicry pattern. The more common mimicry pattern IPHICLUS (forewing with orange subapical spot and white band) is inferred as ancestral, but repeated convergent evolution is also recovered. Evolutionary convergence is also observed for the second most abundant mimicry pattern, COCALA (orange-white banded). Increased rates of mimicry pattern evolution are also found toward the equator. These results underscore the complexity of mimicry evolution in the Neotropical limenitidines i.e., *Adelpha* and *Adelphina*, emphasizing the need to explore its interplay with other biotic and abiotic factors.

## Introduction

The striking patterns of butterfly wings have always puzzled evolutionary biologists and furthered their understanding of the processes driving biological evolution. This is especially true for mimicry, the convergence of colour patterns that warn, or deceive, predators and thereby benefit the potential prey species, and which is demonstrated perhaps more spectacularly among the butterflies than in any other organism. Mimicry has been studied in butterflies as a potential driver of speciation (e.g., Jiggins *et al*., 2006; Mullen, 2006), as colour patterns are subject to natural selection by predators (Mallet & Barton, 1989; Willmott *et al*., 2017; Merril *et al.,* 2012) and sexual selection (Jiggins *et al.,* 2001). Mimicry patterns are also associated with larval host plants (Beccaloni, 1997a; Willmott & Mallet, 2004), microhabitats (De Vries, 2003; Elias *et al.,* 2008), altitude (Chazot *et al*., 2014), climatic niches (Doré *et al.,* 2023), disturbance (Uehara-Prado & Freitas, 2009), flight morphology and behaviour (Srygley 1994, 1999, Hill 2021, Page *et al*. 2024); therefore shifts in wing mimicry patterns may be associated with other ecological shifts that can rapidly drive speciation.

The nymphalid genus *Adelpha* (Limenitidinae) is a species-rich genus (87 species; Prudic *et al*., 2002; Willmott, 2003a, 2003b; Willmott & Hall, 2013; Hui-Yun Tseng *et al*., 2022, moved six species into *Limenitis*), distributed from the temperate western USA to southeastern Brazil, Argentina and Paraguay, and a promising model system to investigate the role of wing pattern evolution in speciation processes. These butterflies show marked changes in dorsal wing colour patterns, both at the intraspecific level and between closely related species (22% of *Adelpha* species present within-species polymorphisms; Willmott, 2003a), while more distant species may show extreme resemblance. There are numerous examples of close correspondence between distribution ranges of *Adelpha* subspecies, where dorsal wing patterns change synchronously from one region to another, such as a complex of species that has a narrow white dorsal band in western Ecuador but a broad white band in the north of Venezuela (Willmott, 2003a). The convergence of warning colour patterns towards those found in *Adelpha* is also observed in more distantly related butterfly groups, such as *Doxocopa* Hübner (Apaturinae), e.g., *Doxocopa linda carwa* Lamas, which occurs sympatrically in western Ecuador with similarly narrow-white-banded *Adelpha* species such as *A. iphiclus* (L.)*, A. iphicleola* (Bates)*, A. erotia erotia* (Hewitson) f. *lerna* (Willmott, 2003a)), *Prepona* Boisduval (Nymphalidae: Charaxinae), and certain Riodinidae (e.g., *Synargis* Hübner). Following Aiello (1984), Neild (1996), Willmott (2003a, 2003b) and Ebel *et al*. (2015), we posit that the convergent similarities in wing patterns in *Adelpha* are likely associated with mimicry. Such mimicry could be Batesian, based on the putative unpalatability of certain *Adelpha* species (e.g., Aiello, 1984; Ebel *et al*., 2015; Finkbeiner et al. 2017, 2018). Batesian mimicry between unpalatable and palatable limenitidine species often occurs in temperate zones (Brower & Brower, 1972; Prudic, Skemp, & Papaj, 2007; Prudic, Shapiro & Clayton, 2002; Ritland and Brower, 1991). There is evidence for unpalatability in a species of *Adelpha* which seems to be a model for *Limenitis lorquini* Boisduval in temperate zones. Feeding responses of avian predators showed that birds usually demonstrated long handling times, feather ruffling and bill wiping after consuming *A. bredowii* Geyer compared to *L. lorquini* (Prudic, Shapiro & Clayton, 2002). However, evidence for unpalatability in Neotropical *Adelpha* is lacking, with experiments showing that putative unpalatable species in *Adelpha* are consumed by avian predators in the field (Carlos E.G. Pinheiro 1996; Robert B Srygley and Chai 1990)(but see Finkbeiner et al. 2017, 2018). An alternative hypothesis for colour pattern convergence in the genus is Müllerian mimicry, based on the ability to escape predators (Mallet & Singer, 1987; Willmott, 2003a; Páez *et al*., 2021). *Adelpha*, with their fast and erratic flight, might be unprofitable to predators because they are difficult to catch (Mallet & Singer, 1987; Willmott, 2003b). *Adelpha* certainly lacks features such as elongate wings, slow flight and a tough body that characterize classically chemically protected aposematic butterflies such as Heliconiini (Brower et al., 1963; Mallet & Singer, 1987; Chai, 1986). As with the evolution of any other trait that might be involved in speciation, it is instructive to examine factors that might affect the evolution of mimicry patterns. For example, it has been hypothesized that the latitudinal diversity gradient (LDG) is linked to stronger interactions among species in the tropics (Schemske *et al*., 2009; Roslin *et al*., 2017). Since dorsal wing patterns in *Adelpha* are potentially involved in signalling both to predators and conspecifics (Dang A., *et al.,* 2025), greater predation pressure or competition for mates in tropical regions could be associated with higher rates of mimicry pattern evolution, i.e., shifting from one mimetic wing colour pattern to another, or the appearance of novel mimetic colour patterns, in such regions. Similarly, tropical lineages are also expected to have shorter generation times, faster mutation rates at higher temperatures (Rohde, 1992), all of which might be expected to drive higher rates of evolution of adaptive phenotypic traits, such as mimicry pattern. Other factors that might accelerate mimicry pattern evolution include abundance and habitat-specialisation. Rare species are likely to be under stronger selection by predators to converge on locally common habitat-generalist species (Birskis-Barros *et al*., 2021). Range-size matters too, as clades containing narrowly distributed habitat-specialist species tend to harbour a higher diversity in mimicry patterns because of adaptation to different, locally abundant mimetic communities (Birskis-Barros *et al*., 2021).

Stronger selection for mimicry in more tropical regions could thus help drive diversification, and previous studies have supported the idea that tropical *Adelpha* clades are diversifying rapidly. *Adelpha* is the main representative of the tribe Limenitidini in the American tropics, along with a few species in *Limenitis* Fabricius, which is widespread in temperate America. Mullen *et al*. (2011) tested the idea that these two clades originated from two independent colonisations of the Americas, with the greater diversity of *Adelpha* being the result of an earlier colonisation and longer time for speciation. A dated molecular phylogeny refuted this hypothesis, and instead supported the hypothesis of a more rapid diversification in *Adelpha* due to multiple host shifts on diverse plant families (Ebel *et al.,* 2015). In particular, Ebel *et al*. (2015) showed that a shift to Rubiaceae may have been a significant event in *Adelpha* evolution, while also confirming the convergent evolution of *Adelpha* dorsal wing patterns.

A more complete understanding of *Adelpha* diversification and its causes requires sorting out remaining systematic challenges. Contrary to the previous hypothesis of Willmott (2003a, 2003b) and Lamas (2004), Mullen *et al*. (2011) did not recover a monophyletic *Adelpha* as conceived by those authors, with a small clade of montane species, known as the *alala-*group, sister to a clade of Palearctic *Limenitis*. This finding has been further supported by subsequent studies (e.g., Dhungel & Wahlberg, 2018; Chazot *et al*., 2021), including a much larger molecular dataset obtained using genome-wide restriction-site-associated sequencing (Ebel *et al*., 2015). In the most recent study of Limenitidinae by Hui-Yun Tseng *et al*. (2022), based on the analysis of mitogenomes and three nuclear ribosomal loci, the *alala*-group was found in a clade grouping the two major *Limenitis* lineages, which led the authors to transfer the members of the *alala*-group to the genus *Limenitis*. However, this taxonomic change based on molecular evidence (mostly mtDNA data) may be premature as it is contradicted by nuclear data and is not supported by specific morphological or ecological evidence; therefore - conservatively - members of the *alala*-group were included in our analysis to attempt to clarify their relationships and because they are co-mimetic with other *Adelpha*.

In the absence of a more comprehensive and robust phylogenetic hypothesis for the genus *Adelpha* and American limenitidines, it is difficult to explore the role of time, colonisation, speciation or ecological interactions such as mimicry on the diversity and distribution of the genus. In the present study, we first attempt to clarify the relationships and classification of *Adelpha* and the *alala-*group, which represent the Neotropical limenitidines, by inferring the most comprehensive phylogeny for these species. Further, we infer ancestral states for wing mimicry patterns and examine the role of shifts in wing patterns in speciation within the Neotropical limenitidines. Finally, we explore various macroecological species traits that might affect its evolution.

## Materials and methods

We assembled a multi-marker molecular dataset, encompassing representatives of 83 of the 87 *Adelpha* species and all six of the *alala-*group species recently placed in *Limenitis* (Hui-Yun Tseng et al., 2022), including 130 subspecies out of 160 subspecies (approx.) (see Table S1). The missing *Adelpha* species are as follows: *A. bredowii*, which is clearly closely related to *A. californica* and *A. eulalia*; *A. gavina*, an endemic southeastern Brazilian species whose relationships are unclear and not strongly supported as related to any particular species or species group, based on the morphological phylogeny of Willmott (2003a); *A. stilesiana*, identified as the sister species to *A. rothschildi* based on morphology; and *A. levona*, which appears closely related to either *A. justina/A. olynthia* based on morphology, or to *A. rothschildi* based on DNA barcoding.

Ten additional species from *Limenitis* (representing 48% of the genus diversity; Mullen, 2006) were included as outgroups: *L. amphyssa* Ménétriés, *L. archippus* Cramer, *L. arthemis* Drury, *L. ciocolatina* Poujade, *L. disjuncta* Leech, *L. glorifica* Fruhstorfer, *L. lorquini* Boisduval, *L. moltrechti* Kardakoff, *L. populi* (L.), and *L. weidemeyerii* Edwards. In addition, two other species from the tribe Limenitidini: *Pandita sinope* Moore and *Parasarpa zayla* Doubleday were included. The tree was rooted with another Limenitidini species, *Moduza urdaneta* Felder & Felder based on the results of Chazot et al. (2021).

### Molecular datasets

We used 771 nucleotide sequences of 16 gene fragments, which were obtained by two different techniques: Sanger sequencing (322 sequences) and RNA-Seq (449 sequences). Our research group generated 580 new sequences and combined those with 191 sequences available on GenBank (see Table S1). For published sequences, we assumed all outgroups were correctly identified. For the ingroup, the great majority of published sequences used were submitted by one or more authors of this paper, who examined voucher specimens, or where the vouchers are figured on BOLD. We also confirmed the identity of samples providing multiple gene sequences by comparing their DNA barcode with a larger barcode library for *Adelpha*, including vouchers that we personally identified.

#### Sanger sequencing data

131 new sequences belonging to 69 *Adelpha* species and four from the *alala*-group were extracted from fresh and museum specimens collected in the last 25 years. Sanger sequences included: the 5’ portion from the mitochondrial genome cytochrome *c* oxidase subunit I (*COI*), and three nuclear gene fragments (all coding): Ribosomal Protein S5 (*RpS5*), glyceraldehydes-3-phosphate dehydrogenase (*GAPDH*) and Elongation factor 1 alpha (*EF-1a*).

#### RNA-seq data

RNA was extracted from multiple tissues using TRIzol and Qiagen kits, following preservation in RNAlater and protocols from Casas et al. (2016). Transcriptome library preparation followed Maytin et al. (2018), using high-quality RNA (RIN ≥ 8), and sequencing was performed on the Illumina HiSeq 2000. Reads were quality-checked (FastQC), trimmed, and normalized using Trinity (Haas et al., 2013). Contigs <500 bp were filtered out with a custom script to improve N50. Transcriptome completeness was assessed with BUSCO, and annotation was performed via BLAST against UniProt, Swiss-Prot, and NCBI NR databases, with GO terms assigned accordingly (Blake et al., 2015).

We used the Sequence Capture Processor (SECAPR) pipeline (http://htmlpreview.github.io/?https://github.com/AntonelliLab/seqcap_processor/blob/master/docs/documentation/main_doc.html) to extract 449 sequences of interest from 48 species (44 *Adelpha,* two *alala-*group and two *Limenitis* species) from the *de novo* annotated sample molecular dataset (see full process in Appendix 1). RNA-seq included 15 nuclear gene fragments (all coding) similar to Dhungel & Wahlberg (2018), whose sequences were available for several *Adelpha* species: Ribosomal Protein S5 (*RpS5)*, Glyceraldehyde-3-phosphate dehydrogenase (*GAPDH),* Elongation Factor 1-alpha *(EF-1a),* carbamoyl phosphate synthetase (*CAD*), Ribosomal Protein S2 (*Rps2*), Arginine Kinase (*ArgKin*), Isocitrate dehydrogenase (*IDH*), dopa-decarboxylase (*DDC*), Cyclin Y (*CycY*), exportin-1-like (*Exp1*), sorting nexin-9-like (*Nex9*), DNA-directed RNA polymerase II polypeptide (*PolII*), suppressor of profiling 2 (*ProSup*), proteasome beta subunit (*PSb*), and UDP glucose6 dehydrogenase (*UDPG6DH*).

#### Specimen-level dataset

The concatenated molecular dataset included 145 taxa for a total length of 19,184 base pairs (bp) made of the following coding gene fragments: *COI* (632 bp), *CAD* (1,335 bp), *RpS5* (381 bp), *Rps2* (816 bp), *GAPDH* (993 bp), *EF-1a* (1,389 bp), *ArgKin*; (1,065 bp), *IDH* (1,230 bp), *DDC* (1,428 bp), *CycY* (1,008 bp), *Exp1* (3,210 bp), *Nex9* (1,617 bp), *PolII* (822 bp), *ProSup* (1,116 bp), *PSb* (696 bp) and *UDPG6DH* (1,446 bp). All sequence datasets were subjected to verification steps by gene fragment using Codoncode Aligner (v3.7.1.1, CodonCode Corporation, http://www.codoncode.com/), through inspection of chromatograms, trimming of low-quality regions, and resolution of mismatches and ambiguities. The list of taxa, GenBank accession codes, and data matrix are available in Table S1.

### Molecular phylogenetic analyses

The phylogenetic analyses were carried out under maximum likelihood (ML) using IQ-TREE v2.2.2.7 (Minh *et al*., 2020). The specimen-level dataset was split into 48 partitions *a priori*, with three partitions (one per codon position) defined for all genes. The Bayesian Information Criterion implemented in IQ-TREE through ModelFinder (Kalyaanamoorthy *et al*., 2017) was used to select best-fit substitution models and partition schemes (see Table S2). Based on the results of the comprehensive study of Chazot *et al*. (2021), *Moduza urdaneta* was used to root the tree. The best-fit ML tree was obtained using 20 independent heuristic searches with the following settings: hill-climbing nearest neighbor interchange (NNI) search (*-allnni* option), partition-resampling strategy (--*sampling GENE* option), best partition scheme allowing the merging of partitions (-*m MFP+MERGE* option) and a perturbation strength of 0.2 (-*pers* 0.2). Branch support for all analyses was assessed using 1,000 replicates for both SH-like approximate likelihood ratio tests (SH-aLRT; Guindon *et al*., 2010) and ultrafast bootstraps (uBV; Minh *et al*., 2013). According to the authors’ recommendations, nodes with SH-aLRT values > 80% and uBV values ≥ 95% were considered highly supported. We also ran ultrafast bootstrap optimization (-*bnni* option) and results showed that original support estimates are robust (see the corresponding .tre files in the Dryad repository for comparison of the resulting trees). Additional analyses were also carried out on individual genes (corresponding outputs are available in the Dryad repository).

### Topological tests

Topological tests were implemented under ML for potential cases where a particular *Adelpha* species was not recovered as monophyletic. To do this, approximately unbiased (AU) tests (Shimodaira, 2002) were carried out with IQ-TREE to determine whether the non-monophyly of the species was supported or not (see Results section). The results of the unconstrained analysis (see above) were compared with results of constrained analyses where non-monophyletic *Adelpha* species were successively constrained to be monophyletic. Apart from these topological constraints, the same exact settings were used for the constrained analyses under IQ-TREE. For the AU tests, the number of resampling of estimated log-likelihoods (RELL) replicates was set at 100,000 (*-zb 100000* option).

### Molecular dating analyses

Further, a subset of the specimen-level dataset was generated in which all *Adelpha* + *alala-*group species were represented by a single individual (preferably one corresponding to the nominate subspecies), with the exception of *A. iphiclus* Linnaeus and *A. iphicleola* Bates for which two additional lineages were kept (see Results section). The resulting species-level dataset (104 taxa) was further used for the molecular dating analyses. Divergence times were estimated using Bayesian relaxed clocks as implemented in BEAST v1.10.4 (Suchard *et al*., 2018). For this study, we implemented a node-dating approach relying on secondary calibrations derived from the study of Chazot *et al*. (2021). The following calibration points were enforced using uniform distributions: *(i)* most recent common ancestor (MRCA) of *Adelpha* + *alala-*group and *Moduza* (minimum age of 22.2 million years ago (Ma) and maximum age of 31.57 Ma); *(ii)* MRCA of *Adelpha* + *alala-*group and *Limenitis* (minimum age of 15.1 Ma and maximum age of 21.49 Ma); *(iii)* MRCA of *Adelpha gelania* Godart and *Adelpha justina* Felder & Felder (minimum age of 12.96 Ma and maximum age of 18.69 Ma); *(iv)* MRCA of *Adelpha nea* Hewitson and *Adelpha paraena* Bates; minimum age of 6.66 Ma and maximum age of 12.59 Ma). Based on the result of the Modelfinder analysis, eight partitions were used and eight distinct uncorrelated lognormal clocks were implemented, and the tree model was set to a birth-death speciation process (Gernhard, 2008). BEAST analyses consisted of four independent runs of 50 million generations of Markov chain Monte Carlo (MCMC) with parameters and trees sampled every 5,000 generations. A 25% burn-in was further applied, and the maximum clade credibility (MCC) tree annotated with median ages and their 95% HPD were produced using TreeAnnotator v1.10.4, which is part of the BEAST software package. Convergence of runs was evaluated graphically and by looking at the effective sample size (ESS) of relevant parameters under Tracer v1.7.2 (Rambaut *et al*., 2018), using the recommended threshold of 200 for relevant parameters.

### Reconstruction of the evolution of mimicry wing colour patterns

#### Mimicry classification

We used a classification of wing colour patterns into mimicry complexes in *Adelpha* + *alala-*group (Willmott, 2003a; Ebel *et al.,* 2015) to code mimicry patterns for all taxa (see Results section). Several studies have used a similar approach for other butterfly groups (Beccaloni, 1997b; Jiggins *et al*., 2006; Doré *et al*., 2021), supported by the fact that birds perceive similarity between species in a similar manner to humans (Dittrich *et al.,* 1993). Additionally, in *Adelpha* + *alala*-group, dorsal wing patterns are simple, few complexes co-occur in each locality, and an expert-opinion-based classification appears sufficient given the typical absence of ambiguous patterns. Fourteen mimicry pattern complexes were recognised (five of which are species-specific), which were used as character states for the species in all subsequent analyses, including within-species polymorphisms: fourteen species present two different colour patterns, four species have three different colour patterns, and two species have four different colour patterns.

#### Ancestral character state reconstructions and phylogenetic signal in Neotropical limenitidines

Ancestral character state reconstructions (ASR) of the wing colour pattern complexes i.e., mimicry patterns, and assessment of their mode of evolution, in particular, whether colour pattern shifts were associated with speciation events (i.e., punctuated evolution) were performed under a Maximum Likelihood framework using BayesTraits v4.0 (Pagel *et al*., 2004). We selected BayesTraits because it allows for the inclusion of polymorphic species and can efficiently handle a large number of discrete character states, which is critical in our case given the 14 mimicry patterns present. First, we estimated the branch scaling parameter kappa defined by Pagel (1999), where branch lengths are raised at the power kappa. A value of one corresponds to gradual evolution (i.e., the probability of colour pattern shift depends on branch lengths), while a value of zero corresponds to a punctuational model of evolution, where trait shift is associated with cladogenesis (i.e., speciation events) rather than branch length, which may indicate that trait shift causes reproductive isolation. Models were compared using AICc scores and the best value of kappa (which was inferred to be equal to 1) was used to infer ancestral states for colour patterns using a maximum likelihood approach. We used the Multistate method, which is suitable for categorical traits that adopt a finite number of states and that allows for polymorphism (Pagel *et al*., 2004). To reduce the number of parameters to estimate, we constrained the probability for all state changes to be equal. The command-lines for the specified nodes to be reconstructed as required by the *addMRCA* and *addNode* commands in BayesTraits were generated with the graphical tree viewer tool BayesTrees v1.4 (https://www.evolution.reading.ac.uk/BayesTrees.html). The ASR was carried out on the BEAST dated phylogeny with *Limenitis* spp. and outgroups pruned. We also performed supplementary ASR analyses using the same phylogeny either on *Adelpha* only (hence *Adelphina* spp., *Limenitis* spp. and outgroups were pruned) or on the clade grouping *Adelpha*, *Adelphina* and *Limenitis* (considering the patterns found in *Limenitis* as a single additional colour pattern).

To test for phylogenetic signal in mimicry pattern evolution, we used the phylogenetic *D*-statistics (Fritz & Purvis, 2010) for binary traits (one state at a time), as implemented in the *phylo.d* function from the *R* package *CAPER* (Orme *et al*., 2013). We ran 1,000 permutations and simulations to test whether observed values of *D* were significantly different from those obtained if we assume a random distribution i.e., no phylogenetic signal (*D = 1),* or under Brownian expectation i.e., phylogenetic signal (*D* = 0). Values from the *D*-statistics were calculated for each mimicry pattern with more than three taxa.

#### Traits associated with rate of wing colour pattern evolution

We tested the power of different species-specific factors to predict rate heterogeneity in wing colour pattern evolution across the tree. For this, we calculated a tip rate of wing colour pattern evolution. We modelled 1,000 histories of colour pattern by sampling at every node in the tree a colour pattern state with a probability equal to that inferred with BayesTraits ASR. For every simulation, we summed the number of transitions from the root to each tip and divided by the number of nodes connecting the root with each tip. The resulting scores were used as tip rate of colour pattern evolution and we calculated for each species the median tip rate out of the 1,000 simulated histories. Mean number of independent origins per colour patterns were extracted from modelled histories of colour patterns. Each transition into a different pattern was counted as a separate event and interpreted as an independent origin of that colour pattern. Tip rate of colour pattern evolution was computed following Chazot *et al.,* (2025)(detailed description in Table S4.1).

Macroecological trait values, used as predictors of colour pattern rate of evolution, were derived by modeling the distribution of all *Adelpha + alala*-group species using locality data compiled and georeferenced through various resources, including Google Earth, published gazetteers, internet searches (e.g. iNaturalist.org), and others.

We computed the following macroecological traits for each species: *(i) tropicality*, measured as the number of degrees from the equator of the geographic distribution centroid, *(ii) density*, measured by the number of specimens (information obtained from multiple public and private international collections [Willmott, 2003], sight records [KW], personal communication [KW], records from publications and web pages) examined per 1,000 km2 of geographic range, *(iii) geographic range size*, and *(iv) niche breadth*, calculated using the product of centered and scaled ranges of suitable values of mean annual temperature, temperature seasonality [standard deviation x 100], mean annual precipitation, and precipitation seasonality (coefficient of variation) across the modeled geographic distribution of each species (chosen for their ability to predict distributions of *Adelpha* species; see Appendix 2, 3 for details). These variables reflect processes that are potentially important to mimicry evolution, including the effect of latitude on diversification rates and biotic interactions (Mittelbach *et al*., 2007; Schemske *et al*., 2009), the influence of relative abundance on selection for mimicry (Müller, 1879), the role of geographic range in color pattern and taxonomic divergence (Endler, 1977), and the importance of climatic niches in mimicry (Chazot *et al*., 2013; Doré *et al*., 2023). Clearly, some of these are relatively crude measures, and in particular *density* is likely influenced by multiple factors, including sampling and detectability bias, but we nevertheless feel that they are likely to capture broad-scale patterns of interest.

To investigate a relationship between mimicry pattern rates of evolution and each predictor described above, we performed Phylogenetic Generalized Linear Models (PGLS) using the *pgls* function (Grafen, 1989; Martins & Hansen, 1997) in the *R* package *CAPER* (Orme *et al.,* 2013). With this function we fitted a linear model accounting for the phylogenetic signal by estimating and applying the branch scaling parameter lambda (Pagel, 1999) using maximum likelihood. Lambda was inferred to be equal to 1 in all analyses.

## Results

### Phylogenetic and dating analyses

The best-fit ML tree from the IQ-TREE analyses of the specimen-level dataset was relatively well-supported overall with *ca*. 63% of the Shimodaira–Hasegawa-like approximate Likelihood Ratio Test (SH-aLRT) values ≥ 80% and *ca*. 78% of the uBV ≥ 95% (see Figure 1). The genus *Adelpha* as historically conceived was found to be polyphyletic and consists of two distinct clades that are highly-supported (SH-aLRT of 99.9% and uBV of 100% for the ‘montane’ *Limenitis* clade (*alala*-group) and SH-aLRT of 99.4% and uBV of 100% for the ‘lowland’ clade). The relationships between these two clades and the two other *Limenitis* clades are not strongly supported (SH-aLRT < 80% and uBV < 95%). In the Taxonomy section below, we describe a new genus for the *alala*-group and discuss why we believe this is the best taxonomic approach. All *Adelpha + alala*-group species for which several subspecies were sampled are recovered as monophyletic with the exception of the following species: *A. erymanthis* Godman & Salvin, *A. hyas* (Doyère), *A. iphicleola, A. iphiclus, A. leuceria* (Druce), *A. lycorias* (Godart), *A. messana* Felder & Felder*, A. plesaure* Hübner, *A. radiata* Fruhstorfer*, A. thessalia* (Felder & Felder). For four out of these 10 species (*A. erymanthis*, *A. iphicleola, A. iphiclus* and *A. leuceria*), the non-monophyly caused by the placement of several subspecies was also statistically supported by the results of the AU-tests (see Table S3), leading us to retain the corresponding subspecies in subsequent dating and ASR analyses. We also propose several taxonomic changes for non-monophyletic species for which we have additional ecological evidence (see the Taxonomy section below).

**Figure 1.**
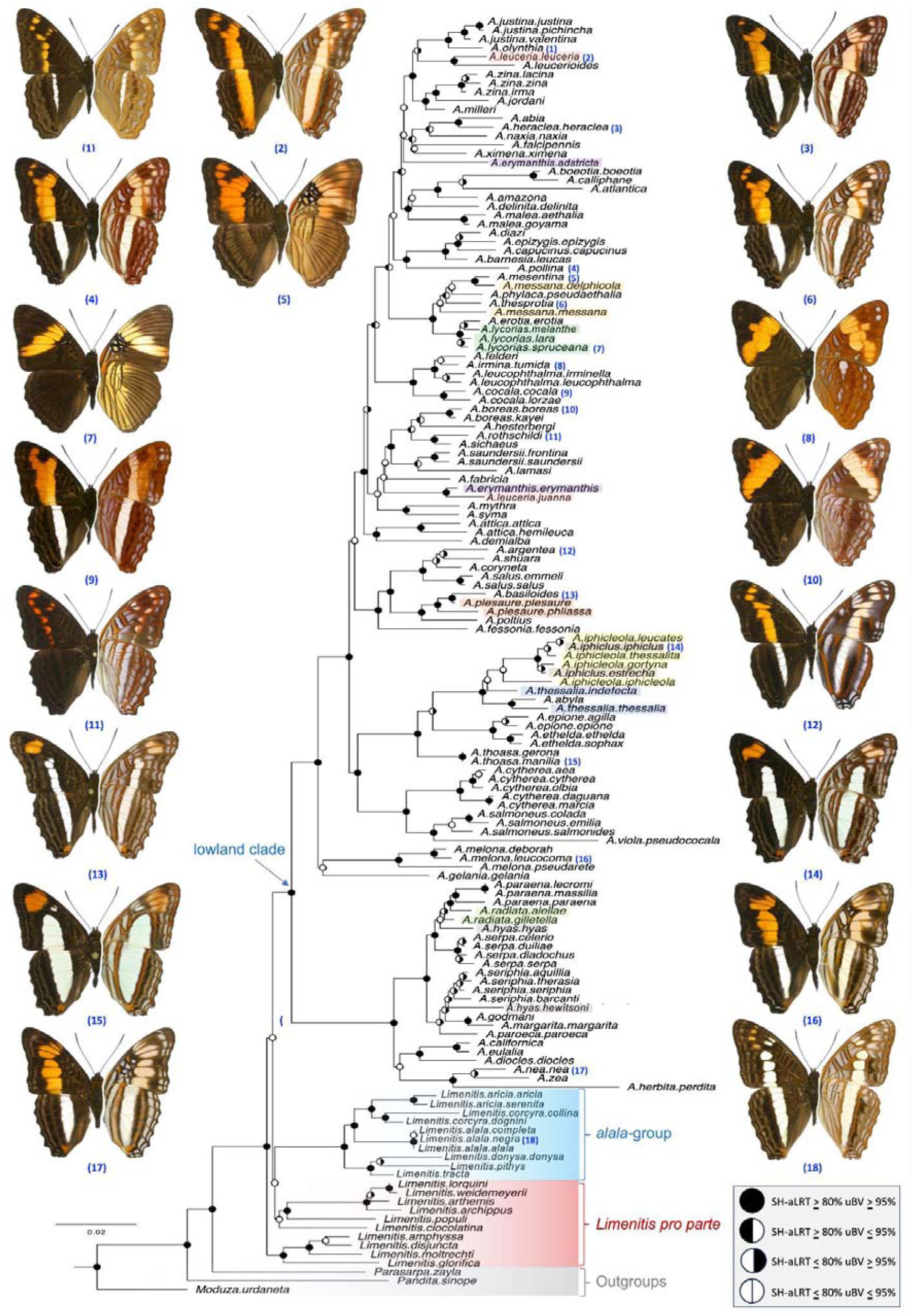
Best-fit ML tree from the IQ-TREE analyses of the specimen-level dataset. Branch support values from phylogenetic analyses (SH-aLRT and uBV, respectively) are represented by semicircles on nodes. The large lowland clade comprising all extant *Adelpha* species is underlined using an arrow. The clade corresponding to the *alala*-group is also highlighted in light blue, whereas remaining *Limenitis* and outgroups are highlighted in red and grey, respectively. Pictures of male adults (dorsal wings on the left, ventral wings on the right) are provided for 17 species; one picture of a female of *A. leuceria* is included as well (all pictures by K. Willmott).

The post-burn-in parameters of the BEAST analyses showed ESS ≥ 200 for all relevant parameters (i.e. those related to age estimates). In the resulting dated phylogeny (see the online supplementary material for the .tre file; see also Figure 4) the ‘lowland’ (*Adelpha*) clade was estimated to have originated during the Middle Miocene *ca*. 16.09 Ma (95% HPD: 18.49-14.15 Ma) while the ‘montane’ (*alala*-group) clade is estimated to have originated during the Late Miocene *ca*. 7.34 Ma (95% HPD: 9.14-5.69 Ma).

**Figure 2.**
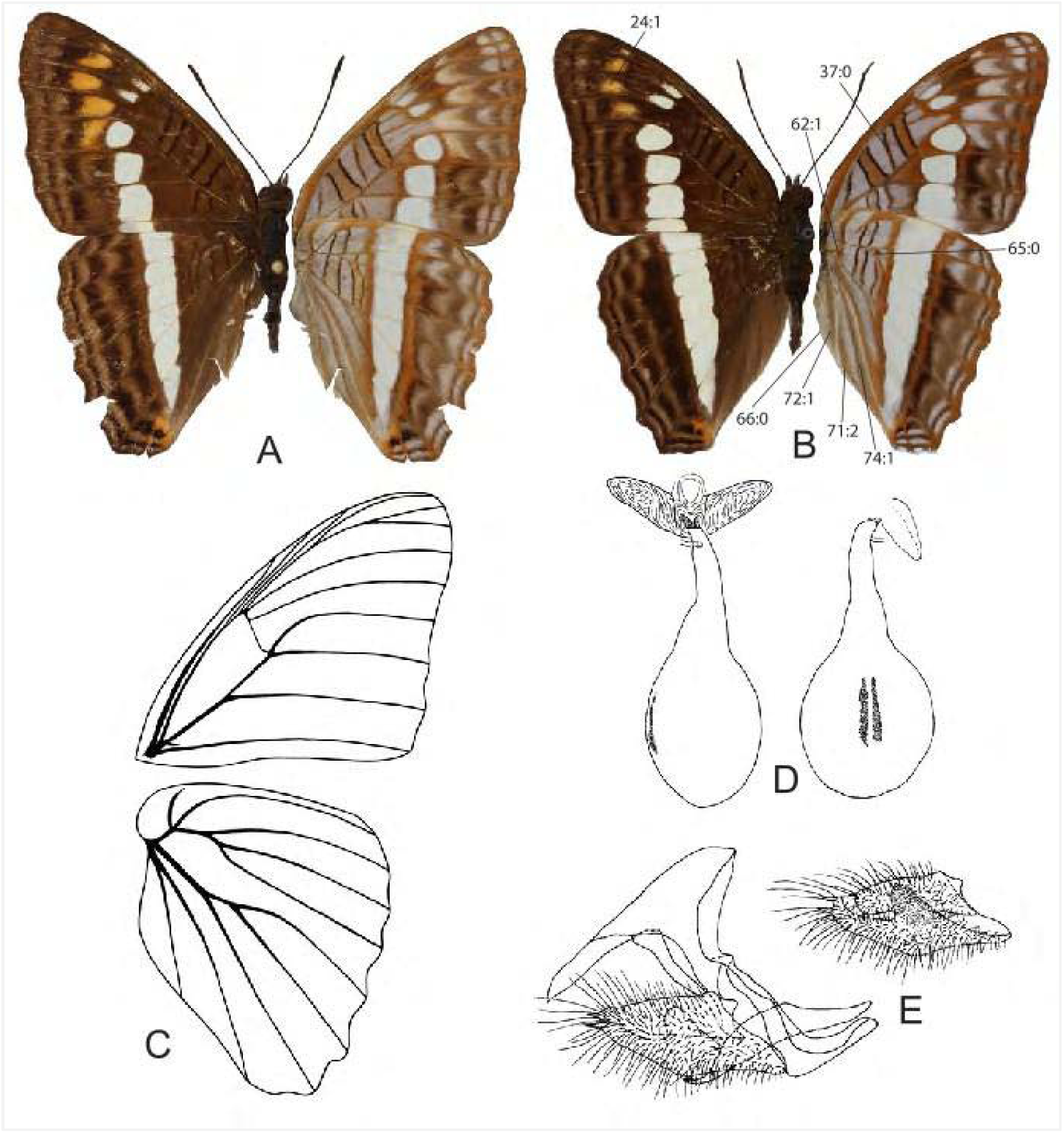
Morphology of *Adelphina* **n. gen. A,B**: Dorsal (left) and ventral (right) wings of female (A) and male (B) *Adelphina alala negra* **n. comb.** from Ecuador. Lines in B indicate characters discussed in the text, numbered as in Willmott (2003a). **C**: Wing venation of *Adelphina alala negra*, Ecuador. **D**: Female genitalia of *Adelphina alala negra*, dorsal view (left), and perpendicular view of signa (right) (from Willmott, 2003a). **E**: Male genitalia of *Adelphina alala negra*, lateral view (left), and inner view of valva showing clunicula (right) (from Willmott, 2003a).

**Figure 3.**
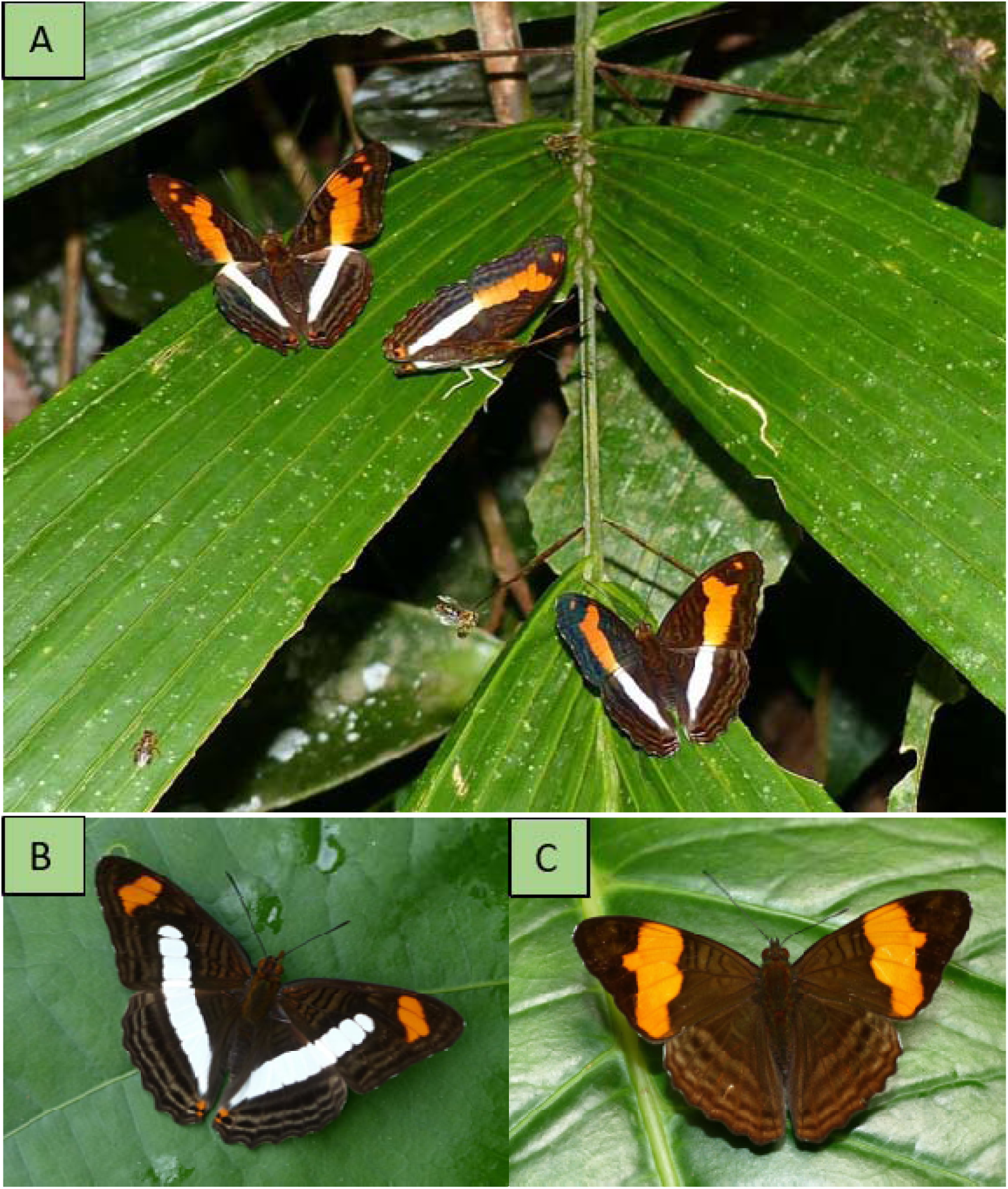
Most widespread mimicry patterns in *Adelpha.* (A) Three sympatric species of *Adelpha* harbouring the mimicry pattern COCALA (from top to bottom): *A. pollina* Fruhstorfer, *A. fabricia* Fruhstorfer, *A. jordani* Fruhstorfer. (B) Most widespread mimicry pattern IPHICLUS (*A. iphiclus*). (C) SALMONEUS mimicry pattern (*A. saundersii* Hewitson). Pictures by Andrew Neild.

**Figure 4.**
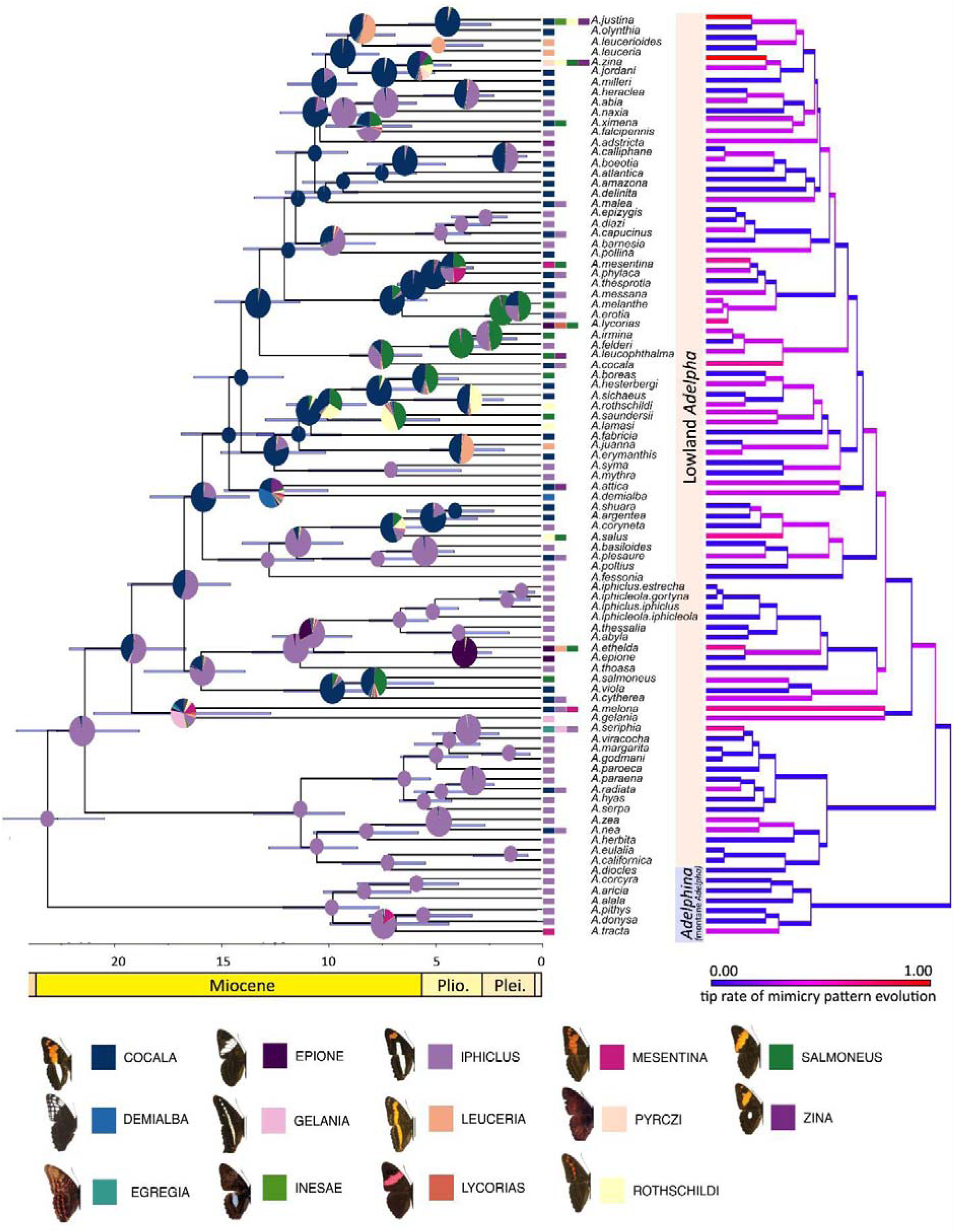
Wing colour pattern evolution. Ancestral character states probabilities, based on the BEAST tree, are shown as pie charts at nodes, using the same colours as terminal squares for species mimicry patterns. Smaller pie charts represent the nodes with more than 0.99 probability of a state. Rates of wing colour pattern evolution are illustrated on coloured branches by a gradient scale, reflecting colour pattern shifts standardized by nodes from root to tip. Species names account for the proposed taxonomic changes. Horizontal bars on nodes represent 95% HPD of age estimates. Outgroup taxa (*Limenitis amphyssa*, *L. archippus*, *L. arthemis*, *L. ciocolatina*, *L. disjuncta*, *L. glorifica*, *L. lorquini*, *L. moltrechti*, *L. populi*, *L. weidemeyerii*, *Pandita sinope*, *Parasarpa zayla*, and *Moduza urdaneta*) were pruned to highlight only the Neotropical Limenitidini and improve visualization of ancestral states within *Adelpha* and *Adelphina*. Wing colour pattern illustrations are adapted from Ebel *et al*. (2015).

### Taxonomy

***Adelphina*** Páez & Willmott, **new genus**

Type species: *Heterochroa alala* Hewitson (1847: 261, Pl. XXI, fig. 8), by present designation.

**Systematic placement and diagnosis.** *Adelphina* **n. gen.** is strongly supported as monophyletic based on molecular data (this study, SH-aLRT of 99.9% and uBV of 100%) and morphological data (Willmott, 2003b; bootstrap 96%, decay index 0.8). In our study based on molecular data, the genus is also strongly supported (SH-aLRT values ≥ 80% and uBV ≥ 95%) as a member of a clade of Limenitidina that contains *Adelpha* and several clades of North American, Asian or Palearctic species, including four North American species (for which the generic name *Basilarchia* could apply), *Limenitis populi* (the type species of *Limenitis*), *Limenitis ciocolatina* (‘*Sinimia*’), and a clade containing *Limenitis amphyssa*, *L. disjuncta*, *L. moltrechti* and *L. glorifica* (‘*Ladoga*’). However, as in previous studies, the relationships among *Adelphina* **n. gen**., *Adelpha s. s.*, and other clades of *Limenitis*, are not strongly resolved.

Within the Limenitidina, *Adelphina* **n. gen.** is diagnosed by possessing a venal stripe on the VHW along vein 3A that is split into anterior and posterior portions either side of the vein, with the anterior portion faded and most pronounced in the distal half (not even throughout or on the vein, or heavy and broken at the base) (Figure 2B; Willmott, 2003b, character 71:2, Fig. 3Y in that paper), and posterior portion reduced to a small dash near the base of the vein (not even throughout or on vein) (Willmott, 2003b, character 72:1, Fig. 3Y). Other species of Limenitidina either lack a venal stripe at vein 3A, or have an even stripe along the vein, or a stripe that is split either side of the vein but not in the configuration found in *Adelphina* **n. gen.**

Characters shared with *Adelpha* that distinguish *Adelphina* **n. gen.** from other limenitidines include: **1)** the postdiscal series on DFW fused to form a subapical marking (see Figure 2B; Willmott, 2003b, character 24:1); **2)** a dark streak at the base of the VHW discal cell (see Figure 2B; Willmott, 2003b, character 62:1); **3)** a dark brown longitudinal line on the VHW from base of cell 3A-2A to middle of anal margin in this cell (see Figure 2B; Willmott, 2003b, character 74:1).

Characters that distinguish *Adelphina* **n. gen.** from *Adelpha* include: **1)** the VHW discal cell with the postcellular bar distinct, continuing approximately parallel to the first cell bar into cell M1-Rs (see Figure 2B; Willmott, 2003b, character 65:0, Fig. 3Y); **2)** the ventral half of the thorax where the legs fold is pale (dark in *Adelpha*, Willmott, 2003b, character 4:0); **3)** the VHW anal margin distal edge is the not lined with dark scales see Figure 2B; Willmott, 2003b, character 66:0).

### Etymology

The generic name *Adelphina* is an arbitrary word based on the name *Adelpha*, intended to indicate a close affinity in geography, wing pattern and historical classification among these species. It is treated as a feminine noun.

### Description

Some notable characters include (numbers in parentheses refer to character states in Willmott (2003b) as indicated on Figure 2B): Labial palpi laterally white, lacking dark central line (1:0); Ventral thorax uniformly pale, lacking dark lines where legs fold (4:0). Dorsal forewing with inner and outer postdiscal series (*sensu* Willmott, 2003b) fused into single orange or white blocks within each cell (24:1). Ventral forewing with third discal cell bar (37:0). Ventral hindwing with: discal cell with dark line at base (62:1); postcellular bar distinct, continuing approximately parallel to the first cell bar into cell M1-Rs (65:0); vein 3A with anterior portion of venal stripe faded and most pronounced in distal half (71:2); posterior portion of venal stripe reduced to a small dash near the base of the vein (72:1). Venation as illustrated (Figure 2C), five FW radial veins with both R1 and R2 originating before discocellulars, spur vein typical of Limenitidini at base of FW cubital vein. Valvae relatively squat with 1-4 short teeth at posterior tip (Figure 2E), clunicula present (Figure 2E) and in some species broad and reduced; corpus bursae with two parallel, elongate signa (Figure 2D). Wings and genitalia of all taxa are illustrated by Willmott (2003a).

### Distribution and natural history

*Adelphina* **n. gen.** contains six species confined to Neotropical montane cloud forest, including three occurring in Central America from Mexico to Panama, and three in the tropical Andes from Venezuela to northern Argentina (Willmott, 2003a, Luis-Martínez et al. 2003, Luis-Martínez et al. 2022). Willmott (2003a) summarized knowledge of the distribution, habitats, behavior and immature stage biology for all species. Adults of *Adelphina* **n. gen.** species appear to be involved in mimicry with one another as well as other *Adelpha*, and the immature stages are notable for feeding on *Viburnum* (Adoxaceae), making leaf shelters, and having greatly reduced body scoli (see Willmott, 2003a for more details). Some immature stage morphological characters may represent synapomorphies for *Adelphina* **n. gen.**, but confirming that possibility requires a detailed comparative study of material from across the Limenitidini.

### Discussion

Based on morphological characters, Willmott (2003b) defined an “*alala*-group” that contained the six species here placed in *Adelphina* **n. gen.**, and found this group to be sister to all other species then placed in *Adelpha*. Willmott (2003b: 300) stated [figure numbers refer to that paper]: “The monophyly of *Adelpha* is strongly indicated in this study, supported by five synapomorphies, including: the orange inner postdiscal series in the dorsal forewing apex (23: 1, Fig. 2V), fusion of the postdiscal series on the dorsal forewing (24: 1, Fig. 2V), the presence of a dark line (basal streak) at the base of the ventral hindwing discal cell (62: 1, Figs 1; 3Y), a dark stripe along vein 3A on the ventral hindwing (68: 1; Figs 1, 3Gg) and a dark stripe in the middle of cell 3A2A on the ventral hindwing (74: 1; Figs 1, 3Y, Z).” Clearly, these are all wing pattern characters, and the lack of genitalic or wing venation characters that typically help to define genera was noted by Willmott (2003b).

A number of molecular phylogenetic studies have since focused on the Limenitidini, with varying conclusions. Mullen et al.’s (2011) study recovered a clade containing all *Adelpha* species along with a number of *Limenitis* species (sometimes placed in other genera, see below) from North America, Asia and the Palearctic, which we refer to here as the *Adelpha* + *Limenitis* (AL) clade. Within the AL clade, there is support for a clade containing North American limenitidines (‘*Basilarchia*’), a group containing *Limenitis amphyssa* and relatives (‘*Ladoga*’), *Adelphina* **n. gen.**, and *Adelpha s. s.* (with the removal of *Adelphina* **n. gen.**). Subsequent molecular studies (Ebel et al., 2015; Dhungel & Wahlberg, 2018; Chazot et al., 2021; Hui-Yun Tseng et al., 2022; this study) have all recovered the same clades, but the relationships among those clades and other key species, such as *Limenitis populi* and *Limenitis ciocolatina*, are consistently only weakly supported and vary across studies. Nevertheless, no molecular study has found support for a monophyletic *Adelpha sensu* Willmott (2003a,b), with *Adelphina* **n. gen.** being placed as sister to remaining AL species excluding *Adelpha s. s.* (Mullen et al., 2010; Ebel et al., 2015), with a polytomy between *Adelphina* **n. gen.**, *Adelpha s. s.* and a clade containing remaining AL species (Dhungel & Wahlberg, 2018), as sister to *L. populi* + *Basilarchia* (Chazot et al., 2021), as sister to *L. ciocolatina* + *L. cleophas* (‘*Sinimia*’) (Hui-Yun Tseng et al., 2022), or as sister to *L. populi* + *Basilarchia + L. ciocolatina* (‘*Sinimia*’) (this study).

To summarize, all phylogenetic studies to date agree that *Adelphina* **n. gen.** and *Adelpha s. s.* are monophyletic, but disagree about the relationships among these lineages and remaining AL species. Hui-Yun Tseng et al. (2022) conducted the most taxonomically complete study of the AL clade to date, based largely on mitochondrial loci, and solved the resulting chaos by recognizing only two genera, *Adelpha s. s.* and *Limenitis* (also containing the *alala*-group). We believe that a more helpful classification is to recognize a new genus for the *alala*-group, and, by implication, to also treat several other lineages within *Limenitis sensu* Hui-Yun Tseng et al. (2022) as distinct genera. Firstly, the monophyly of *Limenitis* as defined by Hui-Yun Tseng et al. (2022) is only weakly supported in all studies, with Chazot et al. (2021), for example, recovering a paraphyletic *Limenitis* (*sensu* Hui-Yun Tseng et al. 2022) with respect to *Adelpha*. Secondly, several of the well-supported lineages within the AL clade constitute biogeographically, morphologically, and ecologically distinct groups whose inferred ages (12.4-13.6 Ma) are well within the range of ages of other nymphalid genera (e.g., Chazot et al., 2019). In the case of *Adelphina* **n. gen.**, its species can be readily recognized by a number of wing and body pattern characters, and they occupy a limited geographic distribution in montane regions of Central America and western South America, far from other non-*Adelpha* limenitidines. Thirdly, generic names are already available for most, or perhaps all, of the AL clades that would need them (see Table S5) (Hemming, 1967). Although revising the generic classification of non-American limenitidines is best done by researchers with specific knowledge of those groups, what seems to us as a reasonable possible classification for species contained in recent molecular studies is provided in Table S5. Fourthly, we believe that a more granular generic-level taxonomy is more helpful for users and likely to be more heuristic, as discussed by Espeland et al. (2023b). Fifthly, we note that even though the relationships of *Adelphina* **n. gen.** to other limenitidines are still unclear, there is strong evidence that this clade is not placed inside *Adelpha s. s.* or any other well-supported, generic-level limenitidine clade, and even if it were to prove sister to *Adelpha*, both generic names would remain valid. Thus, we see this as the most appropriate way to stabilize the future nomenclature of this clade, even as more data are applied to resolving its relationships.

As a final observation, we note that the resulting *Adelpha s. s.* as defined here has limited morphological support, with Willmott (2003b: 300) noting “only a single character supports the monophyly of the remaining clade of *Adelpha* exclusive of the *alala*-group: the possession of dark lines where the legs fold against the thorax (4: 1, Fig. 2G)”. Nevertheless, as noted above, the monophyly of *Adelpha s. s.* has been recovered in all subsequent molecular phylogenetic studies.

**Taxa included in *Adelphina*.**

The following taxa and names are transferred from *Limenitis* (see Hui-Yun Tseng *et al*., 2022), with details of the original descriptions and type specimens provided by Willmott (2003a).

***Adelphina*** Páez & Willmott, **n. gen.**

(“-” denotes a subspecies, “--” a synonym and “---” an unavailable name)

***alala*** (Hewitson, 1847) **n. comb.**

*-completa* Fruhstorfer, 1907

*--titia* Fruhstorfer, 1915

*-negra* (C. Felder & R. Felder, 1862)

*--ehrhardi* Neuburger, 1907

*--albifida* Fruhstorfer, 1907

*--cora* Fruhstorfer, 1907

*--fillo* Fruhstorfer, 1907

*--negrina* Fruhstorfer, 1913

*---praecaria* Fruhstorfer, 1915

*--privigna* Fruhstorfer, 1915

***aricia*** (Hewitson, 1847) **n. comb.**

*-serenita* Fruhstorfer, 1915

*-portunus* Hall, 1938

***corcyra*** (Hewitson, 1847) **n. comb.**

*-aretina* Fruhstorfer, 1907

*-collina* (Hewitson, 1847)

*--epidamna* (C. Felder & R. Felder, 1867)

*-dognini* Willmott, 2003

-*salazari* Willmott, 2003

***tracta*** (Butler, 1872) **n. comb.**

*pithys* (Bates, 1864) **n. comb.**

*--vodena* Fruhstorfer, 1915

***donysa*** (Hewitson, 1847) **n. comb.**

*--roela* (Boisduval, 1870)

*-albifilum* Steinhauser, 1974

### Species-level taxonomic changes in *Adelpha*

In our study, molecular data contributed to the refinement of the species-level classification of *Adelpha*. Based on the results of molecular phylogenetic analyses and additional ecological evidence, we propose the following taxonomic changes.

- *leuceria/juanna*: Samples of *A. leuceria juanna* Grose-Smith did not cluster with samples of *A. leuceria leuceria* (see Figure 1), but instead were sister to *A. erymanthis.* Moreover, one of us (Ichiro Nakamura) collected both *A. leuceria leuceria* and *A. leuceria juanna* in close proximity (and without any signs of intergradation) in the Serranía de Pirre (Panama, Darién), with *A. leuceria* at slightly higher elevations. We therefore restore the species status of *Adelpha juanna* Grose-Smith 1898 **rev. stat.** *Adelpha juanna* thus seems to be a South American replacement for *A. erymanthis,* with wing patterns convergent with *A. ethelda ethelda* Hewintson. Nevertheless, it would be desirable to include additional samples of both *A. erymanthis* and *A. pollina* to confirm these relationships.
- *erymanthis/adstricta*: samples of *A. erymanthis adstricta* Fruhstorfer did not cluster with those of *A. erymanthis erymanthis*, but instead the former taxon formed a clade with a number of Amazonian and southeast Brazilian lowland species (see Figure 1). Furthermore, the recent discovery of the former taxon in Costa Rica by Janzen, Hallwachs and Hill (unpublished data) shows that it is broadly sympatric with *A. erymanthis* throughout Costa Rica and western Panama, at least. Willmott (2003a) retained Fruhstorfer’s (1915) original placement of *adstricta* as a subspecies of *A. erymanthis* based on similarities in the ventral wing pattern, but only three female specimens, all with vague or incorrect locality data, were available for examination at that time. Since then, additional material from Costa Rica and western Ecuador permitted not only clarification of the relationships of the taxon based on molecular data, but also a better understanding of wing pattern variation. As a result of collecting a series of specimens in western Ecuador which show continuous variation in sympatry between typical *adstricta* and *A. erymanthis fortunata* Willmott (also from western Ecuador), we synonymize the latter taxon with the former (**n. syn.**), and raise the former to species as *A. adstricta* Fruhstorfer 1915 **n. stat.**
- *A. hyas/A. viracocha*: Samples of *A. hyas hewitsoni* Willmott & Hall from eastern Ecuador grouped with *A*. *seriphia* (Felder & Felder), far from a single sample of *A. hyas hyas* from southeastern Brazil, which grouped with *A. radiata* (see also Rush et al., 2023). Willmott (2003a) tentatively associated *A. hyas* taxa based on similarities in size and ventral colour pattern, but none of these characters can be considered especially strong in the light of the relationships implied by the molecular data. We therefore regard Brazilian *A. hyas* as a monotypic species and place its two former west Amazonian subspecies as a distinct species, *A. viracocha* Hall 1938 **n. stat.** and *A. viracocha hewitsoni* 1999 **n. stat.** This hypothesis of relationships must also be considered provisional and based just on what seems most biogeographically plausible, given the lack of molecular data for *A. viracocha viracocha*.
- *lycorias/melanthe*: These taxa are part of a clade of very closely related but very phenotypically (wing pattern and wing shape) distinct species. Willmott (2003a) regarded *A. lycorias melanthe* (Bates) as the Central American representative of *A. lycorias*, discounting three specimens labelled from Colombia, which would suggest sympatry with the Colombian *A. lycorias melanippe* Godman & Salvin, as mislabelings. This conclusion seemed reasonable given the presence of numerous examples of mislabeled Colombian specimens in collections, the lack of modern Colombian specimens, and the fact that the taxon is otherwise very common everywhere else within its range. Nevertheless, subsequently at least three reliable records of typical *A. melanthe* occurring in western Colombia have come to light, including a specimen from Tamesis (Antioquia), collected by Bruce Aitken, a specimen from Yanaconas (Valle del Cauca) collected by Haydon Warren-Gash (both pers. comm. to Willmott), and a specimen from Titiribi (Antioquia) photographed by Gabriel Jaramillo Giraldo (https://www.inaturalist.org/observations/9147567). The taxon therefore seems to occur widely, if rarely, throughout western Colombia, in broad sympatry with *A. lycorias melanippe*, and we thus treat it once more as a distinct species, *A. melanthe* (Bates 1864) **n. stat**.

### Evolution of mimicry wing colour pattern

#### Ancestral character state reconstructions and phylogenetic signal

When inferring the evolution of mimicry patterns, the best-fit model was obtained for kappa=1, which is more in support of gradual evolution (see Table 1).

**Table 1.**
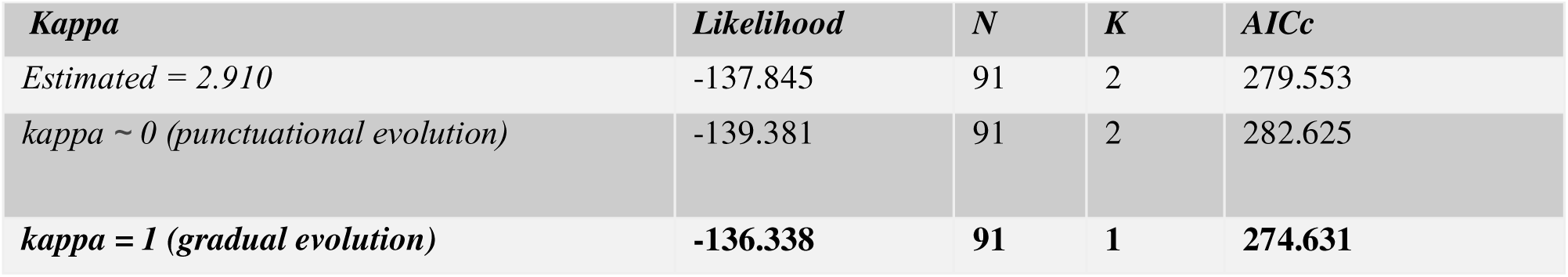
Testing for a punctuational versus gradual mode of trait evolution for the evolution of mimicry pattern. Abbreviations: number of putative species (N), Kappa (K), Akaike’s information criterion with correction for small sample size (AICc). The best model is highlighted in bold.

| *Kappa* | *Likelihood* | *N* | *K* | *AICc* |
| --- | --- | --- | --- | --- |
| *Estimated = 2.910* | -137.845 | 91 | 2 | 279.553 |
| *kappa ~ 0 (punctuational evolution)* | -139.381 | 91 | 2 | 282.625 |
| ***kappa = 1 (gradual evolution)*** | **-136.338** | **91** | **1** | **274.631** |

Estimation of ancestral character states for mimicry colour patterns in *Adelpha + Adelphina* i.e. Neotropical limenitidines showed that IPHICLUS, a colour pattern shared by 47 species and widely distributed across the phylogeny, was most likely to be the state of the common ancestor of all Neotropical limenitidines species (probability = 0.993) (see Figure 3, Figure 4). The reconstruction of colour pattern changes showed mostly unambiguous character states on internal branches. Multiple shifts in wing colour patterns occurred mostly in the lowland clade. Nine mimicry patterns showed multiple independent origins across the phylogeny (see Table S4.2), with COCALA (20.64 times, sd=2.10), IPHICLUS (17.22 times, sd=1.83), and SALMONEUS (11.65 times, sd=0.93) exhibiting the highest numbers of independent origins. The results of the two supplementary analyses (Figs. S1 and S2) consistently supported the result of this analysis, indicating that the crown of both clades are inferred as mostly IPHICLUS.

The phylogenetic signal analysis of mimicry patterns (shared by more than three taxa) based on the *D*-statistics (Fritz & Purvis, 2010) found phylogenetic signals consistent with Brownian motion for IPHICLUS and LEUCERIA which are highly conserved. COCALA, SALMONEUS and ROTHSCHILDI showed a random distribution across the tree, indicating no phylogenetic signal (see Table 2).

**Table 2.**
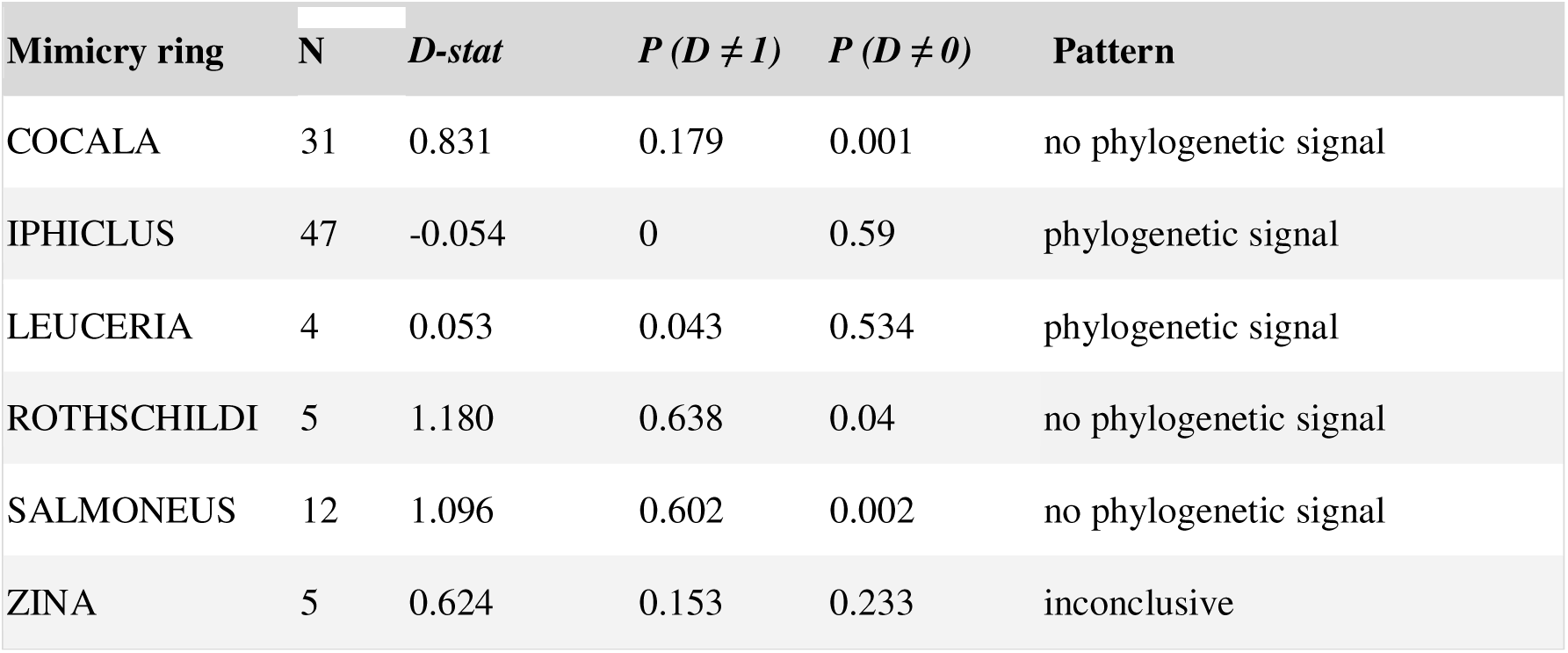
Results of the test for phylogenetic signal in wing colour pattern evolution for each mimicry ring with more than three taxa. N = number of species within each mimicry ring. *D*-stat = *D* score for each mimicry ring (*D* = 1 when a trait is randomly distributed across the tree and *D* = 0 when the trait is distributed according to a Brownian model of evolution). P (*D* ≠ 1) = probability that *D* is significantly different from 1. P (*D* ≠ 0) = probability that *D* is significantly different from 0.

| Mimicry ring | N | *D*-stat | *P* (*D* ≠ 1) | *P* (*D* ≠ 0) | Pattern |
| --- | --- | --- | --- | --- | --- |
| COCALA | 31 | 0.831 | 0.179 | 0.001 | no phylogenetic signal |
| IPHICLUS | 47 | -0.054 | 0 | 0.59 | phylogenetic signal |
| LEUCERIA | 4 | 0.053 | 0.043 | 0.534 | phylogenetic signal |
| ROTHSCHILDI | 5 | 1.180 | 0.638 | 0.04 | no phylogenetic signal |
| SALMONEUS | 12 | 1.096 | 0.602 | 0.002 | no phylogenetic signal |
| ZINA | 5 | 0.624 | 0.153 | 0.233 | inconclusive |

#### Traits associated with rate of wing colour pattern evolution

Inferred species wing colour pattern evolution rates were heterogeneous across the phylogeny (see Figure 4, Table S4.1), and they were significantly correlated with the measure of tropicality with higher rates towards the equator (Figure 5, Table 3). No significant correlations were detected between mimicry evolution and other traits.

**Figure 5.**
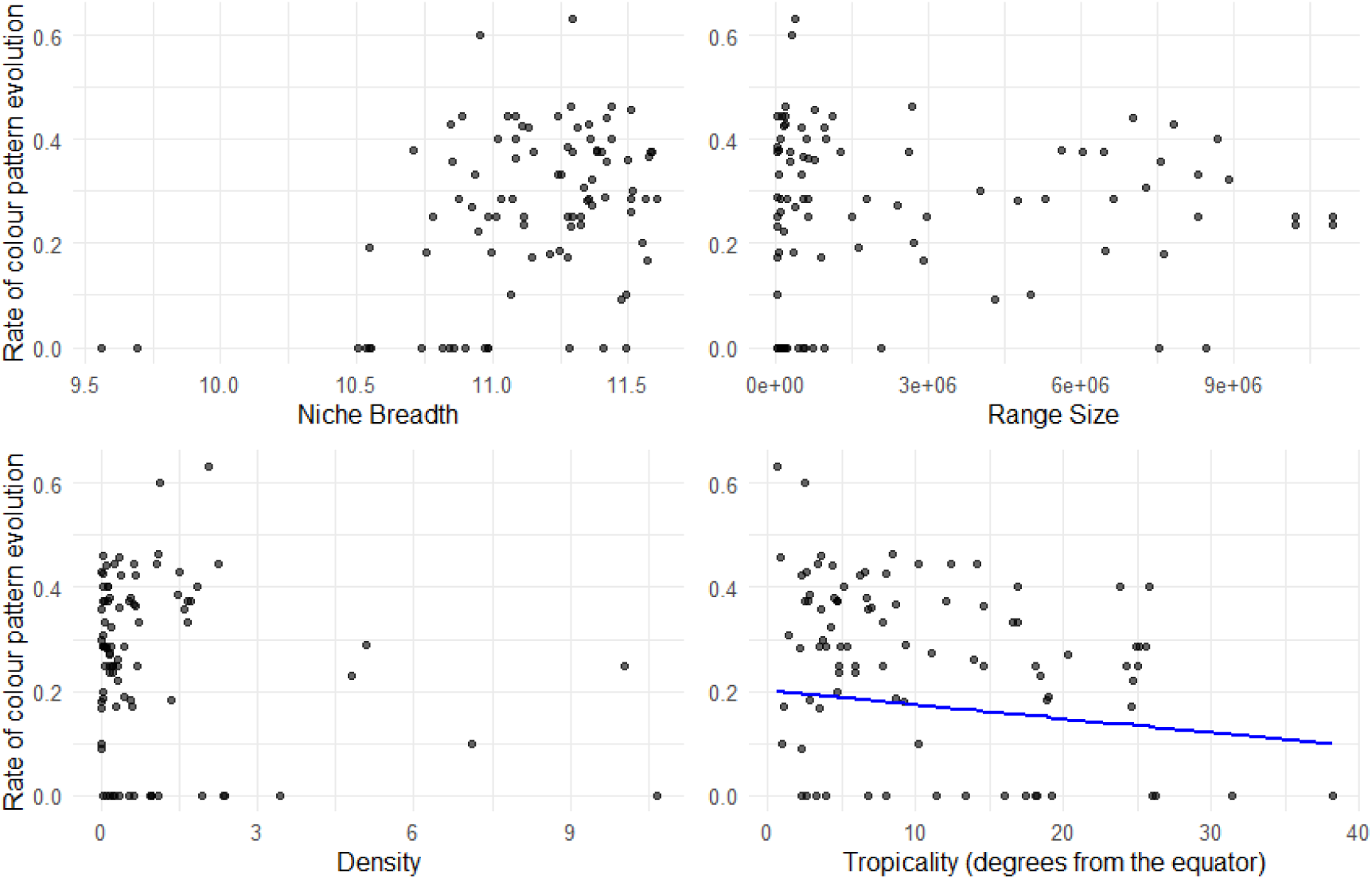
Effect of ecological species traits on wing colour pattern evolution rates. Plots illustrating the relationship between the rate of colour pattern evolution and various species traits: niche breadth (log-transformed), range size, density, and tropicality (measured as degrees from the equator). Among these traits, only tropicality showed a significant association (df = 89, r² = 0.075, p = 0.005). The blue line represents the best-fit regression line from the Phylogenetic Generalized Least Squares (PGLS) analysis.

**Table 3.** Correlates for rates of mimicry pattern evolution and species traits. *Niche breadth value was log transformed

| Predictor | df | Estimate | $r^2$ (adjusted) | t | lambda | p-value |
| --- | --- | --- | --- | --- | --- | --- |
| *Niche breadth\** | 89 | 0.0242 | -0.004 | 0.813 | 1.000 | 0.4181 |
| *Range size* | 89 | 1.0308e-9 | -0.001 | 0.352 | 1.000 | 0.7254 |
| *Density* | 89 | -0.00046 | -0.011 | -0.119 | 1.000 | 0.9056 |
| *Tropicality (degrees from the equator)* | 89 | -0.00265 | 0.075 | -2.886 | 0.998 | 0.0049 |

## Discussion

### Systematics of the genus *Adelpha*

We inferred the most taxonomically comprehensive dated phylogeny for the Neotropical genus *Adelpha* and its six former species recently transferred to *Limenitis* (*alala*-group). We found the montane *alala*-group to be sister with a low support (SH-aLRT of 76.9% and uBV of 70%) to a clade of Eurasian and North American *Limenitis*, rendering *Adelpha* as historically conceived polyphyletic, similarly to Mullen *et al*. (2011) and Ebel *et al*. (2015). Other studies have found the montane *alala*-group in an unresolved polytomy between that clade, remaining *Adelpha* species, and temperate *Limenitis* species (Dhungel & Wahlberg, 2018), embedded in a paraphyletic *Limenitis* (Chazot *et al*., 2021), or embedded deep within a monophyletic *Limenitis* (Hui-Yun Tseng *et al*., 2022) (see Figure S3). The montane *alala-*group is distinctive in comparison with *Adelpha* in their larvae feeding on Caprifoliaceae (Dipsacales) and making leaf shelters, like some *Limenitis* species (Willmott, 2003a; Ebel *et al.,* 2015). A closer relationship between the *alala*-group and *Limenitis* than between the *alala*-group and remaining *Adelpha* certainly seems reasonable. Although Hui-Yun Tseng *et al*. (2022) solved the polyphyly of *Adelpha* as historically conceived by moving the *alala*-group into *Limenitis*, we discuss above why we feel this is not the best solution, and consequently we described a new genus, *Adelphina*, for the *alala-*group.

### Mimicry wing colour pattern evolution

Obtaining a taxonomically comprehensive phylogeny allowed us to explore patterns of diversification. In this particular case, we investigated mimicry pattern evolution and its relationship to speciation and species traits.

#### Ancestral character state reconstructions and the role of shifts of mimetic colour pattern in Neotropical limenitidines speciation

We found that the IPHICLUS mimicry pattern was most likely to be the ancestral state for all Neotropical limenitidines i.e., *Adelpha* and *Adelphina* species, and that it also reappeared several times (17 events of independent origins across the phylogeny), indicating a very dynamic pattern of wing colour pattern evolution. It is the most common pattern in the genus (47 species share this pattern). Otherwise, the COCALA mimicry pattern is inferred to be the ancestral state across most of the large, species-rich lowland clade, and most mimicry pattern shifts occurred later in this clade. Our results are in accordance with the general pattern inferred with a reduced taxon sampling by Ebel *et al*. (2015). Mimetic wing colour pattern is an example of a trait for which ecological selection driven by predation can lead to divergence, with reproductive isolation and speciation as a side effect (Mallet *et al*., 1998; Jiggins *et al.,* 2001, 2006; Jiggins 2008; Chamberlain *et al.,* 2009; Merrill *et al.,* 2011). These types of traits are often coined as ‘magic traits’, and reproductive isolation occurs via assortative mating, which implies both premating isolation (Kronforst *et al.,* 2006; Mavárez *et al.,* 2006; Jiggins *et al.,* 2008; Chamberlain *et al.,* 2009) and postmating isolation as a result of increased predation on non-mimetic, rare hybrids (Mallet & Barton, 1989; Pinheiro, 2003; Arias *et al.,* 2016). However, contrary to our expectations, there is no significant association between shifts in mimicry patterns and speciation in the Neotropical limenitidines, a result supported by the observation that groups of related species often have similar wing colour patterns (e.g., *A. fessonia* and *A. basiloides* [Dang et al. 2025]). In other groups of mimetic butterflies, there are also examples where cladogenesis is not accompanied by a shift in mimetic colour pattern, i.e., where pairs of sympatric, closely related species are near-perfect mimics of each other (e.g., in *Heliconius* Kluk butterflies, Giraldo *et al*., 2008; Jiggins, 2008; Mérot *et al*., 2013). It could be that speciation initially occurred through divergence in factors unrelated to mimicry patterns, such as habitat choice at a finer spatial scale, providing the mechanism for strong premating isolation leading to ecological speciation (Jiggins, 2008). Another possibility is that mimicry patterns in Neotropical limenitidines may well have evolved from adaptive introgression between sympatric populations rather than from common ancestry (Mavárez *et al.,* 2006; Jiggins *et al.,* 2008; Pardo-Diaz *et al.,* 2012; Edelman *et al.,* 2019; Kozak *et al.,* 2021; Thawornwattana *et al.,* 2022). Some *Heliconius* species may be the result of very recent mimetic convergences between hybridising species, possibly through adaptive introgression, rather than speciation without changes in colour pattern (The *Heliconius* Genome Consortium, 2012, Rosser *et al*., 2024). In such cases, in response to selection against reproductive interference, other cues may drive reproductive isolation, such as chemical communication i.e., pheromones (Jiggins, 2008; Mérot *et al.,* 2015). Indeed, it has been suggested that closely related mimetic butterflies not only rely on visual cues but also on olfactory cues for sexual attraction (Poulton, 1907; Boppre, 1978; Vane-Wright & Boppre, 1993).

In *Adelpha*, almost nothing is known about mate recognition (but see Dang et al. 2025), but the extreme resemblance between many species suggests that wing pattern may play only an initial role. Indeed, there are possible hybrid specimens between closely related but phenotypically distinct species, e.g., *A. mesentina* Cramer and *A. thesprotia* Felder & Felder or *A. cocala* Cramer and *A. irmina* Doubleday. Instead, Willmott (2003a) hypothesized that mate recognition and courtship in the genus may be partly mediated by pheromones, noting that *Adelpha* males have a dense area of darker scales at the base of the ventral forewing which is lacking in females and that could represent androconial scales.

It is also possible that speciation in *Adelpha* may have been primarily driven by other ecological factors, such as host plant use. Mullen *et al*. (2011) suggested that the increase in species richness of (lowland) *Adelpha* compared to the montane *alala-group* i.e., now described as the *Adelphina* genus, might be due to adaptive divergence resulting from host plant shifts to Rubiaceae and other families. Additionally, Ebel *et al*. (2015) found phylogenetic evidence of multiple host plant shifts in *Adelpha*, suggesting its possible contribution to rapid adaptive diversification.

Studies on Lepidoptera have shown the correlation between multiple ecological shifts such as forest structure, flight height, host plants and mimicry patterns (Beccaloni, 1997a, DeVries *et al*., 1999; Willmott & Mallet, 2004; Jiggins *et al.,* 2006; Elias *et al.,* 2008; Hill, 2010; Chazot *et al.,* 2014; Ortiz-Acevedo *et al.,* 2020; Dell’Aglio *et al*., 2022), which, in combination or independently, can also lead to speciation. Abiotic factors, such as geography or climate, are likely linked to host plant shifts accompanying speciation as well (Lisa De-Silva *et al*., 2017; Condamine *et al*., 2018; Kergoat *et al*., 2018; Allio *et al*., 2021; Aduse-Poku *et al*., 2022; Espeland *et al*., 2023a; Hévin *et al.,* 2024). Further research should consider the geographical context in host plant - Neotropical limenitidines’ interactions to investigate whether adaptations to new host plants represent post-speciation events after geographic isolation, rather than the main driver of speciation (e.g., Barraclough *et al*., 1999; Jousselin *et al.,* 2013; Doorenweerd *et al*., 2015; Berry *et al*. 2018; Jousselin & Elias, 2019).

In the Neotropical limenitidines, there is very little evidence of links between microhabitats, host plants and mimicry patterns (but see Ebel *et al.,* 2015), and there are few cases of dimorphic species within populations in species harbouring the two most abundant mimicry patterns i.e., IPHICLUS and COCALA (e.g., *Adelpha erotia, A. capucinus*). This might suggest that in *Adelpha* and *Adelphina*, these two major mimicry patterns are maintained by other processes rather than by ecological differences. Mallet’s interpretation of the shifting balance hypothesis (Wright, S., 1977; Mallet & Singer, 1987) may explain polymorphism in the Neotropical limenitidines’ species, where colour patterns that are more or less equally fit may become established due to locally relaxed selection (i.e., little pressure for convergence). Although this remains speculative, other aspects such as mate choice and predation on non-mimetic rare hybrids need to be investigated. Shifts on colour patterns in *Adelpha* and *Adelphina* do not appear to be the main driver of speciation, but this does not rule out a long-term role in generating intraspecific diversity.

#### Phylogenetic signal and convergence in mimetic colour patterns

In the case of the Neotropical limenitidines i.e., *Adelpha* and *Adelphina*, it has been hypothesised that convergence in colour patterns is mainly due to mimicry (Willmott, 2003a, 2003b; Ebel *et al.,* 2015), but closer examination of the phylogenetic signal showed heterogeneous modes of evolution among mimicry colour patterns. The most species-rich mimicry pattern in this group, namely IPHICLUS (47 species), is composed of co-mimetic species that are in general closely related, with a strong phylogenetic signal suggesting that many cases of similarity result from common ancestry rather than convergence. This is most likely explained by IPHICLUS being the ancestral pattern, so it is not surprising that it is present in several communities and lineages (both closely and less closely phylogenetically related). The next most abundant mimicry rings, COCALA (31 species) and SALMONEUS (12 species), showed no phylogenetic signal, thus indicating a higher signal of evolutionary convergence. Our results suggest that evolutionary convergence is important in wing colour pattern evolution in some Neotropical limenitidines lineages, and more likely in others that could not be tested because of the level of representation (e.g., MESENTINA, GELANIA mimicry patterns appear repeatedly in phylogenetically distantly related species in our phylogeny). Additionally, mimicry with the Neotropical limenitidines has also been evidenced in other more distantly related genera such as *Prepona* (Nymphalidae: Charaxinae), *Doxocopa* (Nymphalidae: Apaturinae) and even in one species of *Synargis* Hübner (Riodinidae) (Willmott, 2003a).

Referring to the tribe Ithomiini, Chazot *et al*. (2025) suggested that patterns that appeared earlier could allow for higher accumulation of species, either through speciation without colour pattern shift or through phylogenetic convergence of the mimicry pattern. Moreover, different colour patterns may be associated with different predator selection pressures (Birskis-Barros et al. 2021), e.g., degrees of generalisation, and therefore lead to different rates of convergence and conservatism. Furthermore, the maintenance of a strong phylogenetic signal in mimicry patterns could occur when a clade diversifies within a single area where butterflies are exposed to the same predator community, and where selection in favour of mimicry pattern stability is therefore expected. Further work should focus on the spatial dynamics of diversification as well, which may provide interesting insights on the maintenance of the strong phylogenetic signal in *Adelpha* mimicry patterns.

#### Rate of mimicry pattern evolution increases with tropicality

We found evidence that the rate of mimicry pattern evolution increases at lower latitudes and that this is not mainly due to common ancestry between species, providing novel evidence for the existence of a latitudinal gradient in mimicry evolution. A number of authors have suggested that there may be latitudinal gradients in biotic interactions (Wallace, 1878; Dobzhansky, 1950; Schemske, 2002, 2009), such as higher rates of herbivory (Coley & Aide, 1991) and insect predation (Janzen, 1970; Novotny *et al*., 2006; Roslin *et al*., 2017; Zvereva *et al*., 2019), as well as mutualistic interactions, in the tropics (Schemske *et al.,* 2009). Schemske (2009) has suggested that strong biotic interactions in the tropics promote coevolution, and that, when interacting species coevolve, the optimum phenotype is constantly changing, which could lead to faster adaptation. Ricklefs & O’Rourke (1975) observed a greater variety of sizes, patterns, and shapes in tropical species than in temperate moth species (but see Ricklefs, 2009). Therefore, for *Adelpha*, greater predator diversity and abundance (hence greater complexity in predator-prey interactions), greater inter-/intra-specific competition for mates, and stronger spatial structuring of mimicry patterns, among others, could lead to (or allow) more shifts of mimicry patterns in more tropical species. Although we found no evidence that shifts in mimicry patterns are frequently associated with speciation across the entire genus, there appears to be connections between mimicry patterns and elevation or geographic region. These connections may have facilitated at least some shifts to new elevations (e.g., in *A. jordani* & *A. zina* Hewitson) or new regions (e.g., in *A. sichaeus* Butler, *A*. *rothschildi* Fruhstorfer and *A. levona*) in tropical areas, potentially contributing to the remarkable diversity found in these regions.

## CONCLUSION

The study of mimicry in *Adelpha* makes it an appealing model system for addressing questions regarding the evolution of wing colour pattern diversity, speciation and diversification in Neotropical butterflies. We present here the most taxonomically comprehensive phylogeny for the genus *Adelpha* and related Neotropical limenitidines, which supports the hypothesis that *Adelpha* as historically conceived is not monophyletic. To stabilize future nomenclature we describe a new genus, *Adelphina* **n. gen.**, for the *alala-*group, transferring those species from where they were recently placed in *Limenitis*. We found that mimicry pattern shifts do not appear to be the main driver of speciation in *Adelpha*, since a gradualist model of evolution of mimicry patterns is more likely than a punctuational model. We also found adaptive convergence to be important in the evolution of colour patterns, but also a strong phylogenetic signal in the most represented mimicry pattern, IPHICLUS. Finally, the rate of mimicry pattern evolution is correlated with the tropicality of species, supporting the hypothesis of stronger biotic interactions towards the equator, which may have contributed to the generation and maintenance of more diverse tropical communities. Collectively, our results encourage further research into other factors that could help explain patterns of diversity and diversification in this group of butterflies.

## Author contributions

ERIKA PAEZ V.: Conceptualization; methodology; validation; formal analysis; data curation; investigation; resources; writing – original draft; writing – review and editing; visualization. GAEL. J. KERGOAT: Conceptualization; methodology; validation; formal analysis; data curation; writing – original draft; writing – review and editing; visualization.

NICOLAS CHAZOT: Conceptualization; methodology; validation; formal analysis; writing – review and editing.

MOHAMED BENMESBAH: data gathering.

ADRIANA D. BRISCOE: data gathering, writing - review and editing.

SUSAN FINKBEINER: data gathering, review.

ANDRE V. L. FREITAS: data gathering, writing - review and editing.

ROBERT P. GURALNICK: Conceptualization; methodology; writing - review and editing.

RYAN I. HILL: data gathering, writing - review and editing.

MARCUS KRONFROST: data gathering, review and editing.

LUIZA M. MAGALDI: data gathering, writing - review and editing.

SEAN P. MULLEN: data gathering, writing - review and editing.

ICHIRO NAKAMURA: data gathering - review.

HANNAH OWENS: Conceptualization; methodology; data gathering; formal analysis; writing - review and editing.

NIKLAS WAHLBERG: data gathering, writing - review and editing.

MAXWELL WOODBURY: Conceptualization; methodology; data gathering; formal analysis; writing - review and editing.

MARIANNE ELIAS: Conceptualization; methodology; validation; formal analysis; writing - original draft; writing - review and editing.

KEITH R. WILLMOTT: Conceptualization; methodology; validation; data gathering; writing - original draft; writing - review and editing.

## Acknowledgments

We want to thank the editor Emmanuel Toussaint and three anonymous reviewers for numerous remarks and constructive comments that helped improve this manuscript.

Specimens from Ecuador were collected and exported under the project “Study of the Genetic Diversity and Evolution of the Lepidoptera of Ecuador”, most recent permit number MAATE-DBI-CM-2023-0298, issued by the Ministerio del Ambiente, Agua y Transición Ecologica with the collaboration of the Instituto Nacional de Biodiversidad (INABIO). Specimens from Brazil were collected under permits 10438-8 and 10802-38, issued by Instituto Chico Mendes de Conservação da Biodiversidade (ICMBio), and specimens from Costa Rica under permits R-003-2016-OT-CONAGEBIO and R-021-2016-OT-CONAGEBIO, issued by Comision Nacional para la Gestion de la Biodiversidad (CONAGEBIO). Field work in Ecuador was funded in part by the Leverhulme Trust, the Darwin Initiative, the FLMNH Museum Associates, the National Geographic Society (Research and Exploration Grant # 5751-96) and the US National Science Foundation (# 0103746, #0639977, #0639861, #0847582, #1256742, DEB-1342759). EPV thanks Senescyt (fellowship Programa Becas “Universidades de Excelencia”). AVLF thanks the Brazilian Research Council - CNPq (fellowships 421248/2017-3 and 304291/2020-0), and the FAPESP (grant 2021/03868-8). This study is registered under the Brazilian SISGEN (AD427E3). LMM thanks the Brazilian Research Council - CNPq (grants 151166/2023-4), and the São Paulo Research Foundation (grant 2023/10376-0). Fieldwork in Costa Rica was funded by the US National Science Foundation (DEB-1342706) and University of the Pacific, and was supported by the Organization for Tropical Studies and MINAE. ADB thanks Aline Rangel-Olguin and Aide Macias-Muñoz for assistance with sequencing. We thank Andrew Neild for the authorization to use his pictures, and Laura Braga, Richard Raby, Roberto Greve and Augusto Rosa for collecting some of the Brazilian specimens. We also thank all those who helped collect specimens in the field, contributed to digitizing *Adelpha* specimens in collections, or otherwise provided data.

## Data availability statement

The data that support the findings of this study are openly available in Dryad repository DOI number 10.5061/dryad.7h44j104b

Newly generated sequences were deposited in GenBank, and are registered with the following accession numbers: XXXXXX-XXXXXXX.

