## Supplementary figures and images for "Phylogeny, systematics and evolution of mimicry patterns in Neotropical limenitidine butterflies"

### Figure S1

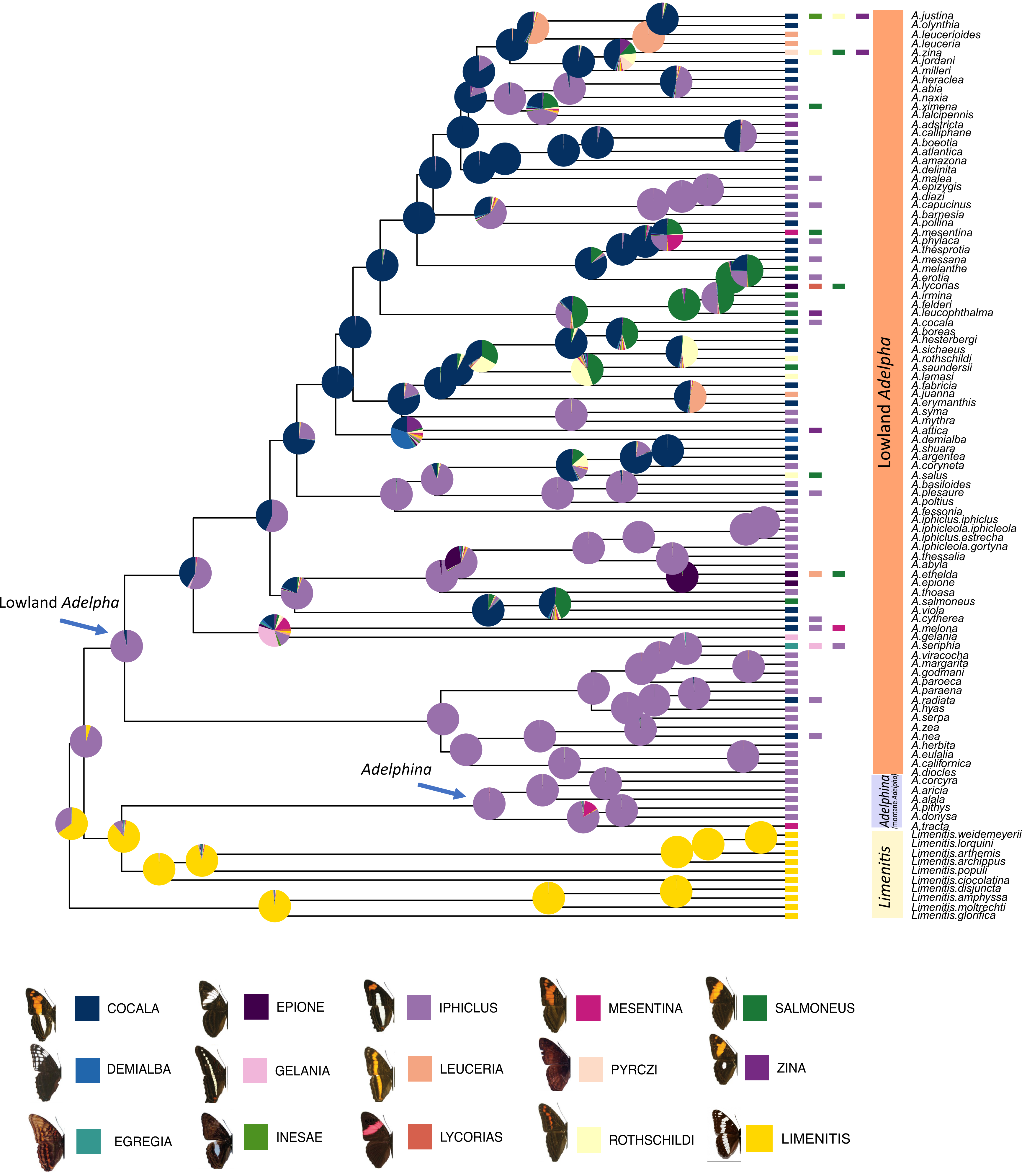

### Figure S2

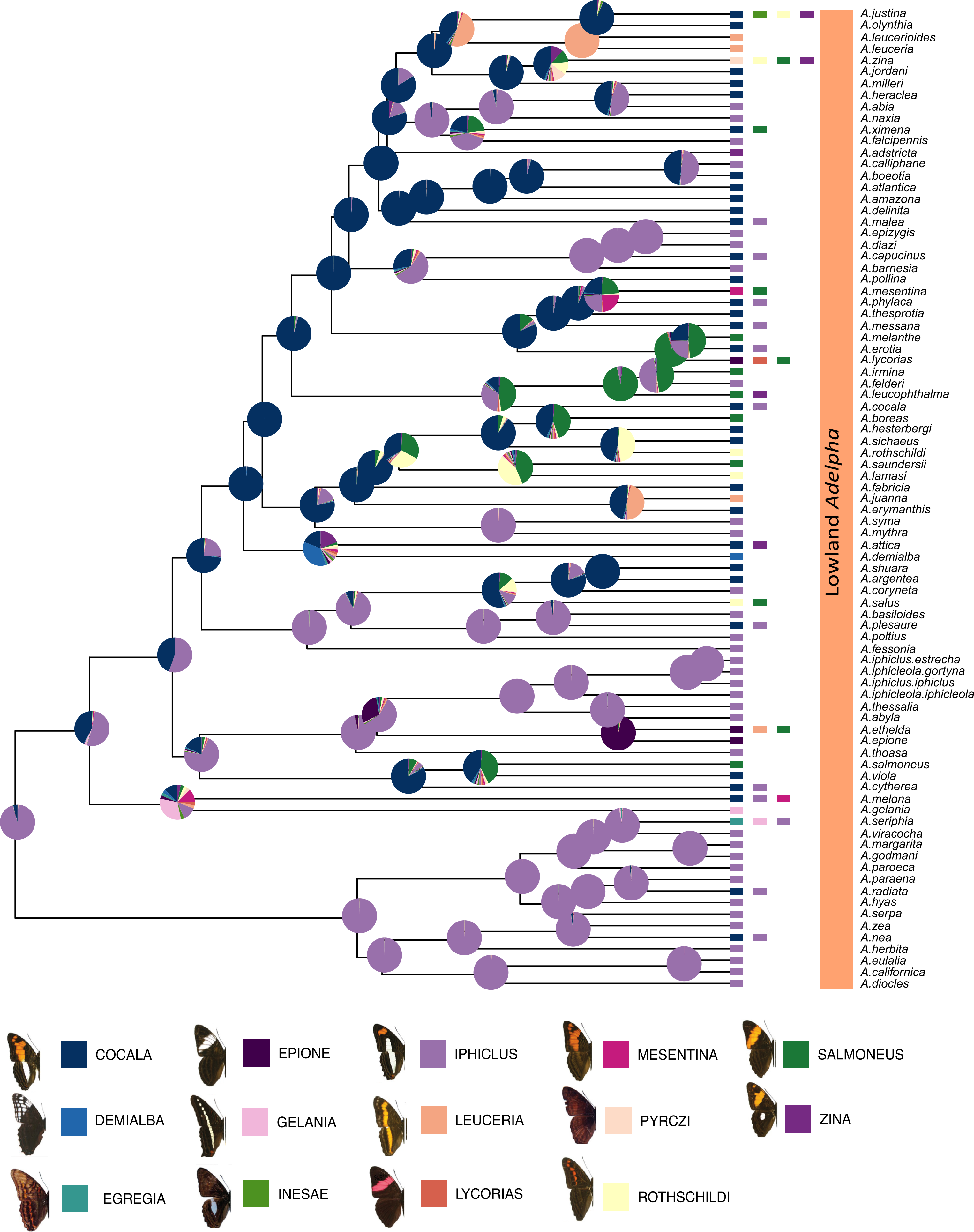
