## Supplementary material for "Phylogeny, systematics and evolution of mimicry patterns in Neotropical limenitidine butterflies": Figure S3

**Figure S1.** A comparison of tree topologies from recent molecular phylogenetic studies, simplified to clades relevant to generic classification.

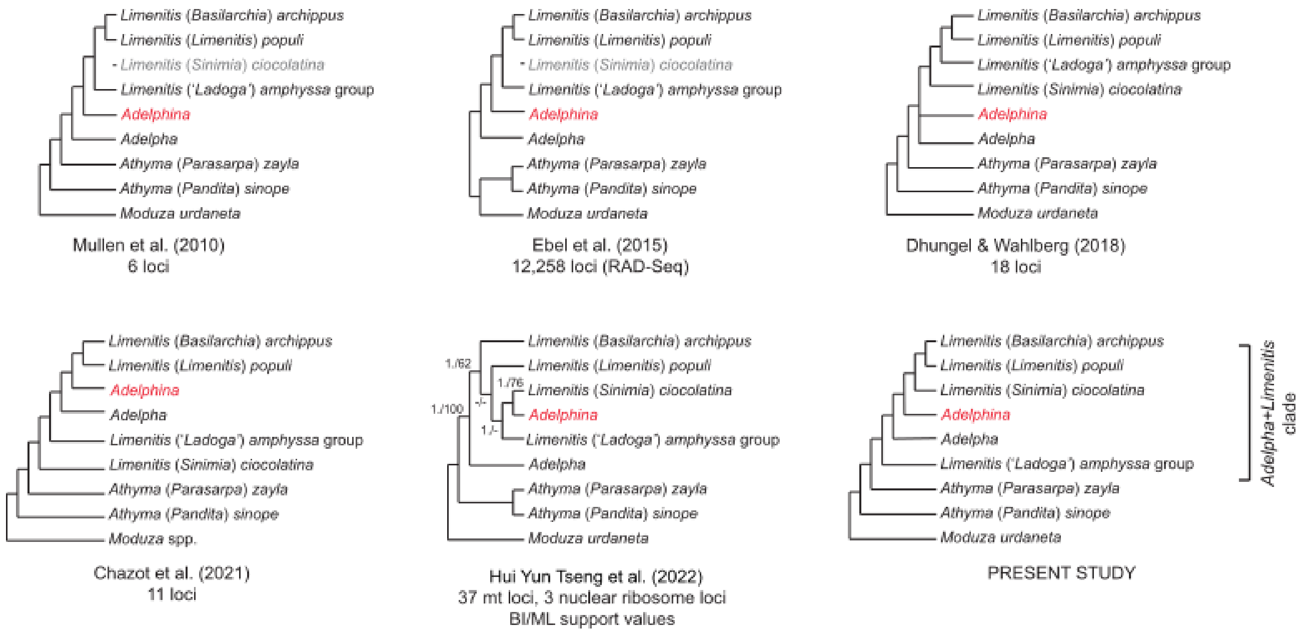
