## Supplementary material for "Phylogeny, systematics and evolution of mimicry patterns in Neotropical limenitidine butterflies": Table S3

| **Table S3.** Results of the AU-tests carried out with IQ-TREE. The constraints that generate trees that are significantly less-supported by the AU-tests (-) are highlighted using a bold font. | | | |
| --- | --- | --- | --- |
| **Constrained tree** | **logL** | **deltaL** | ***p*-AU** |
| **#1: *A. erymanthis* monophyletic** | **-88502.83864** | **20.14** | **0.0182 (-)** |
| #2: *A. hyas* monophyletic | -88498.84976 | 16.151 | 0.0702 (+) |
| **#3: *A. iphicleola* monophyletic** | **-88512.36033** | **29.661** | **0.00372 (-)** |
| **#4: *A. iphiclus* monophyletic** | **-88596.98369** | **114.28** | **7.35e-15 (-)** |
| **#5: *A. leuceria* monophyletic** | **-88570.94941** | **88.25** | **2.98e-42 (-)** |
| #6: *A. lycorias* monophyletic | -88488.59690 | 5.8978 | 0.384 (+) |
| #7: *A. messana* monophyletic | -88483.56740 | 0.86826 | 0.632 (+) |
| #8: *A. plesaure* monophyletic | -88499.47261 | 16.773 | 0.113 (+) |
| #9: *A. thessalia* monophyletic | -88494.51715 | 11.818 | 0.218 (+) |
| #10: *A. radiata* monophyletic | -88486.25171 | 3.5526 | 0.419 (+) |
